# 3D computational models explain muscle activation patterns and energetic functions of internal structures in fish swimming

**DOI:** 10.1101/556126

**Authors:** Tingyu Ming, Bowen Jin, Jialei Song, Haoxiang Luo, Ruxu Du, Yang Ding

## Abstract

How muscles are used is a key to understanding the internal driving of fish swimming. However, the underlying mechanisms of some features of the muscle activation patterns and their differential appearance on different species are still obscure. In this study, we explain the muscle activation patterns by using 3D computational fluid dynamics models coupled to the motion of fish with prescribed deformation and examining the torque and power required along the fish body with two primary swimming modes. We find that the torque required by the hydrodynamic forces and body inertia exhibits a wave pattern that travels faster than the curvature wave in both anguilliform and carangiform swimmers, which can explain the traveling wave speeds of the muscle activations. Most interestingly, intermittent negative power (i.e., power delivered by the fluid to the body) on the posterior part, along with a timely transfer of torque and energy by tendons, explains the decrease of the duration of muscle activation towards the tail. The torque contribution from the body elasticity further solves the mystery of the wave speed increase or the reverse of the wave direction of the muscle activation on the posterior part of a carangiform swimmer. For anguilliform swimmers, the absence of the changes mentioned above in the muscle activation on the posterior part is in line with our torque prediction and the absence of long tendons from experimental observations. These results provide novel insights into the function of muscles and tendons as an integrative part of the internal driving system, especially from an energy perspective, and highlight the differences in the internal driving systems between the two primary swimming modes.

**Author summary:** For undulatory swimming, fish form posteriorly traveling waves of body bending by activating their muscles sequentially along the body. However, experimental observations have showed that the muscle activation wave does not simply match the bending wave. Researchers have previously computed the torque required for muscles along the body based on classic hydrodynamic theories and explained the higher wave speed of the muscle activation compared to the curvature wave. However, the origins of other features of the muscle activation pattern and their variation among different species are still obscure after decades of research. In this study, we use 3D computational fluid dynamics models to compute the spatiotemporal distributions of both the torque and power required for eel-like and mackerel-like swimming. By examining both the torque and power patterns and considering the energy transfer, storage, and release by tendons and body viscoelasticity, we can explain not only the features and variations in the muscle activation patterns as observed from fish experiments but also how tendons and body elasticity save energy. We provide a mechanical picture in which the body shape, body movement, muscles, tendons, and body elasticity of a mackerel (or similar) orchestrate to make swimming efficient.

## Introduction

In the undulatory swimming of fish, a backward-traveling wave of body bending is formed to push against the water and generate propulsion. Muscle is the executor of the neural control and the source of mechanical power in fish swimming. Therefore, how muscles are used is a key question in understanding the control and mechanics of fish swimming and has been under multidisciplinary research over the past decades.

Experimentally, muscle activation during swimming is measured using electromyography (EMG) for various fish species [1–5](Fig 1). During steady swimming, a common pattern emerges: the muscle elements are activated in the manner of a wave traveling posteriorly, but this EMG wave travels faster than the curvature wave [6]. As such, the phase difference between the curvature and EMG varies along the body, known as “neuromechanical phase lags”. Nonetheless, details of the muscle activation pattern vary among species. For anguilliform swimmers such as an eel, the speed difference is not large, and the duration of the muscle activation on one side of the body is approximately half of the undulation period [3]. For carangiform swimmers such as a carp, the propagation speed of the EMG onset is much higher than that of the curvature wave, whereas that of EMG termination is even higher, resulting in decrease of duration towards the tail [4]. The EMG activity, together with muscle contraction kinetics, the strain and the volume of the active muscle, can determine the absolute muscle power output along the body. With this approach, Rome *et al.* [5] showed that for scup, the power is generated mostly by the posterior part of the body.

**Fig 1.**
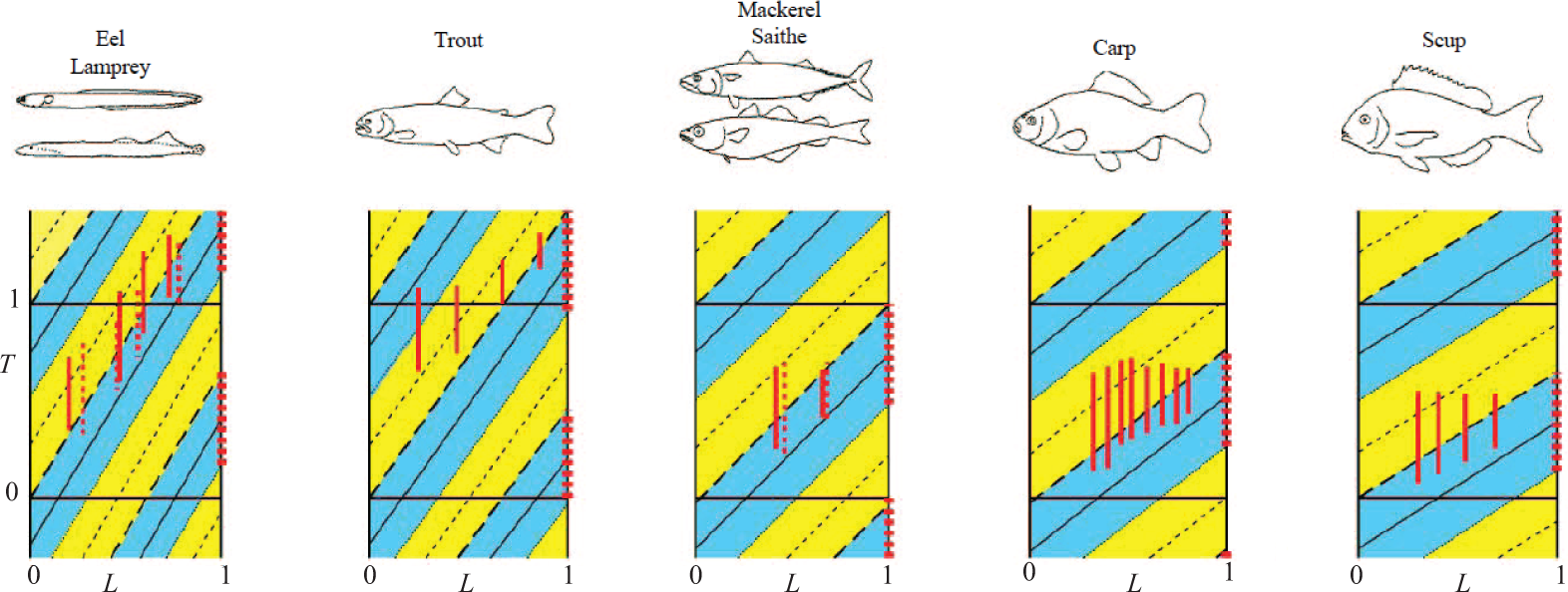
Electromyography patterns for fish species with different body forms and kinematics for steady swimming. The red vertical lines indicate the duration of EMGs of the muscle on the left-hand side at that position on the body (the EMGs of lamprey and saithe are dashed). The yellow and blue stripes indicate the durations when the local curvature is increasing and decreasing, respectively. The red dotted lines on the right side of the figure indicate the duration when the tail is moving from rightmost to leftmost. Adapted from [6] with permission from the Journal of Experimental Biology.

To understand the muscle activation patterns and underlying mechanical principles of internal driving, researchers previously studied the internal torque and the corresponding power required. The sign of the torque has been used to predict which side of the muscle should be activated. Using theoretical models, namely resistive force theory [7], elongated body theory [8], and 3D waving plate theory [9, 10], previous studies obtained torque waves that travel faster than the curvature waves and qualitatively explained the neuromechanical phase lag. However, since positive and negative torques both occupy half of the period all along the body, the decrease of the EMG duration in carangiform swimmers remains an obscure phenomenon.

Another approach used to understand the internal driving in the coupled system is to use neural control signals as an input and observe the kinematics emerging from the coupling of internal driving, the body, and the external fluid. Using resistive force theory and 2D computational fluid dynamics (CFD) with a prescribed uniform muscle activation, McMillen *et al.* [11] and Tytell *et al.* [12] studied lamprey-like swimmers and showed that the same muscle forces can generate body bending with different wavelengths, corresponding to varying magnitudes of the neuromechanical phase lags, depending on passive body properties such as stiffness. However, since the kinematics emerge from the coupling of many components, this kind of approach may generate kinematics that do not match the experimental observations; therefore, the approach may create difficulties in systematically studying the features the muscle activation and in explaining the differences in the muscle activation between species.

These previous modeling studies were all based on either theoretical models with strong assumptions or 2D CFD models, which cannot capture 3D flow around the top and bottom of the fish body and the jet left behind and 3D shapes for carangiform swimmers [13]. Therefore, while qualitative explanations from the models are reasonable, the errors in these predictions are hard to estimate.

To study the features of muscle activations among different species and reveal the underlying mechanical principles, here we use 3D CFD simulations to investigate the torque patterns and power output patterns for a typical anguilliform swimmer and a typical carangiform swimmer. Combining the simulation results with experimental observations, we aim to explain the features and their variations in the EMG patterns among fish with different swimming modes.

## Model and Numerical Methods

Treating water as an incompressible viscous fluid and the fish as moving bodies with prescribed deformations, we developed numerical 3D models of an eel and a mackerel (see Fig 2).

**Fig 2.**
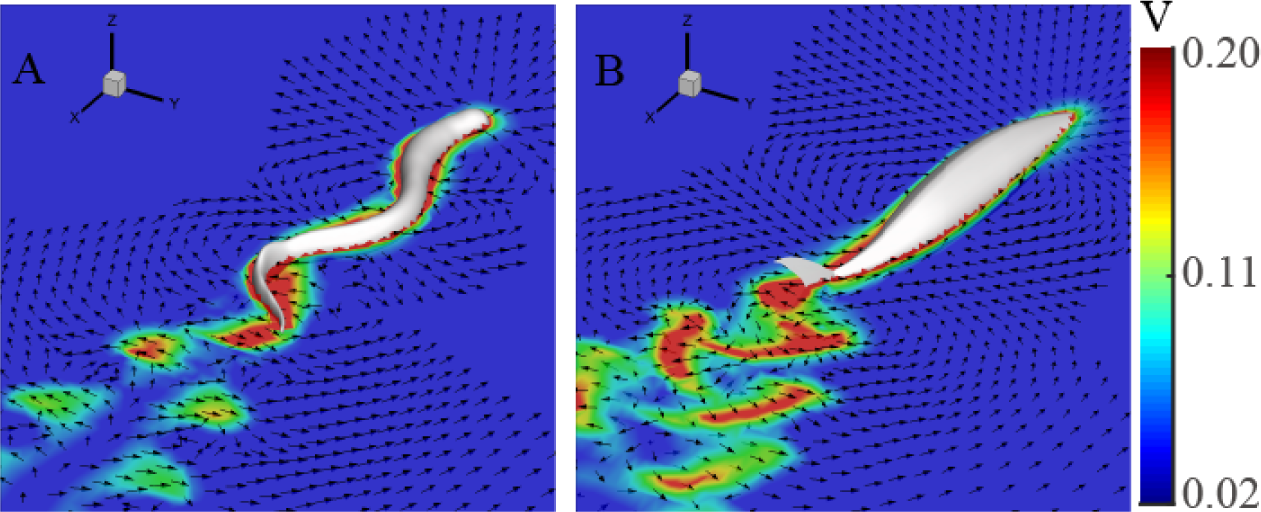
Flow fields in the middle coronal (*z* = 0) planes of eel (A) and mackerel (B) models in the laboratory frame. The arrows represent the velocity direction and the colors represent the magnitude of the velocity. Only flow speeds greater than 0.02 in nondimensionalized units are shown.

### Body shape and kinematics

The carangiform body is modeled based on the actual anatomy of a mackerel, whereas the anguilliform body is created from a lamprey computed tomography (CT) scan (see [14] for details). Except for the caudal fin, other fins are neglected for the swimmers. The lengths of the fish bodies (*L*) are used as the unit length in the simulations. The bodies are meshed with triangular elements, and some sharp and small structures from the scan are removed to avoid an instability of the CFD computation. After obtaining the surface data of the two fishes, we reshaped the fishes and re-meshed the surface grid so that our code could accommodate the boundary between the fish and the fluid. The sharp and thin tail of the mackerel was modeled as a zero-thickness membranous structure. The number of surface mesh points was 3962 for the eel and 2127 for the mackerel (including 1962 for the mackerel’s body and 165 for the tail). See S1 Fig for mesh details. The body mass (*M*) was computed by assuming a uniform distribution of density equal to the fluid density and was 1 in nondimensional units. *M* = 0.0019 for the eel and *M* = 0.0101 for the mackerel.

The kinematics for undulatory locomotion are generally in the form of a posteriorly traveling wave with the largest wave amplitude at the tail. To describe the deformation of the fish bodies, centerline curvatures *κ* are prescribed in the form of *κ*(*s, t*) = *A*(*s*) sin(*ks - ω_u_t*), where *s* is the arc length measured along the fish axis from the tip of the fish head, *A*(*s*) is the amplitude envelope of curvature as a function of *s*, *k* is the wavenumber of the body undulations that corresponds to wavelength *λ*, and *ω_u_* is angular frequency. We use the undulation period as the unit of time, so *ω_u_* = 2*π*. The amplitude envelope *A*(*s*) for the anguilliform kinematics has the form *A*(*s*) = *a*_max_*e^s−^*^1^, where *a*_max_ is the tail-beat amplitude. For carangiform kinematics, the amplitude envelope has the form *A*(*s*) = *a*_0_ + *a*_1_*s* + *a*_2_*s*^2^. The parameters for *A*(*s*) were adjusted to fit the envelope of the movement of real fish observed in experiments [8, 15]. The parameters used were *a*_max_ = 11.41, and *k* = 2*π/*0.59 for the anguilliform swimmer and *a*_0_ = 1, *a*_1_ = -3.2, *a*_2_ = 5.6, and *k* = 2*π/*1.0 for the carangiform swimmer. To avoid generating spurious forces and torques in the interaction between the fish bodies and fluid, we added rotation and translation in the body frame of the swimmers to ensure that the movement of the bodies without external forces satisfied two conservation laws: linear momentum conservation and angular momentum conservation (see S1 Appendix for details). The resulted kinematics are shown in S2 Fig.

### CFD and the fluid-structure interaction

The in-house immersed boundary method code used is capable of simulating 3D incompressible, unsteady, and viscous flows in a domain with complex embedded objects including zero-thickness membranes and general 3D bodies [16, 17]. The flow is computed on a nonuniform Cartesian grid in *x′y′z′* coordinates. The fluid domain has a size of 8.5 × 5 × 5, and a total of 620 × 400 × 400 ≈ 99 million points are used. The grid is locally refined near the body with the finest spacing at 0.005 × 0.005 × 0.005. The fish models are placed in the center of the computation domain, and the body centerlines are in the *z′* = 0 plane. A homogeneous Neumann boundary condition is used for the pressure at all boundaries. The flow speed of the inlet flow and outlet flow at the front and back boundaries are set as the swimming speed of the trial runs so that the model swimmers move only minimally in the computational domain. A zero-gradient boundary condition is used at all other boundaries.

We considered the fluid-structure interaction in the plane of undulation. Because the body of the swimmers is deforming, the governing equation for the angular degree of freedom is d(*Iω*)*/*d*t* = *T*_tot_, where *I* is the moment of inertia, *ω* is the angular speed of the body, and *T*_tot_ is the total torque from hydrodynamic forces. Since the deformation is prescribed, *I* and 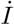 are known. Therefore, *ω* can be obtained by numerically integrating 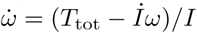 while integrating other equations for the translational movement of the body and the flow of the fluid. The time interval for the integration is 5 × 10^*−*4^.

We set an initial swimming speed of 0.3 at the beginning of the simulation and waited two full cycles for the swimmer to reach steady swimming. All the data presented are collected from the third and fourth cycles. Since the swimming direction is not perfectly aligned with the *x′*-axis of the computation grid, a new coordinate system is used so that the swimming direction is aligned with -*x*, *y* is the lateral direction, and the *z*-axis is the vertical direction. Vertical motion is neglected, but the force magnitude in the *z* direction is only 9.3 × 10^*−*6^ for the eel and 9.3 × 10^*−*5^ for the mackerel, which are less than 3% of the force magnitude in the forward direction. The Reynolds number is defined as *Re* = *U L/ν_k_*, where *U* is the swimming speed, and *ν_k_* = 1*/*15000 is the kinematic viscosity.

### Force, torque, and power in the simulation

The force, internal torque, and power distributions along the fish body as a function of time are computed from the simulation. The force per unit length on the fish body, **F**, is calculated as follows: Take an arc length ∆*s* along the body centerline, and integrate all forces from every mesh point in ∆*s*; then, divide the total force by the arc length ∆*s*.

Considering the hydrodynamic forces, we compute the internal torque required to overcome the hydrodynamic forces and body inertia. The body elasticity and the other internal resistive forces are initially ignored and will be be discussed later. The torque can be computed by integrating the contributions from either side of the body from the point of interest: 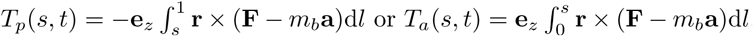, where *m_b_* is the body mass per unit length, and 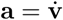 is the acceleration of the body segment. To minimize the numerical error, we use a weighted average of the torques computed from both sides, namely *T* = *sT_p_* + (1 − *s*)*T_a_*.

The internal power by the torque and the power transferred to the fluid per unit length are computed as 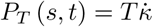 and *P_F_* (*s, t*) = − **F · v**, respectively, where ** is the time derivative of curvature, and **v** is the velocity of the body segment. The difference between the total power computed by integrating the internal power or the external power along the body is within the numerical error (*<* 5%).

## Results and Discussion

### Body movement and fluid flow

The free swimming speeds (*U*) are 0.29 and 0.25 in nondimensionalized units for the eel and the mackerel, respectively. The corresponding Strouhal numbers are 0.63 and 0.68. These values are consistent with previous numerical studies at similar Reynolds numbers (*Re* ≈ 4000) (e.g., [14]). For both fishes, double row vortices are shed behind the tail, similar to previous numerical results (see Fig 2). The velocity field behind the mackerel clearly shows a backward flow, while a mean flow behind the eel in the fore-aft direction is not easily detected.

### Force

As expected from the input kinematics and body shapes, the forces are relatively uniformly distributed on the eel but concentrated on the tail of the mackerel (Fig 3, Fig 4A & C, S1 Video, and S2 Video). The fore-aft and lateral forces both show posteriorly traveling wave patterns similar to those of body bending, except at the head where the surface orientation rapidly changes. For the eel, the peaks in the force components near 0.7 body length correspond to an increase in the body height (in *z* direction) at that position. For the mackerel, the separation of the thrust and drag is clear: the tail generates most of the thrust, and the anterior part of the body generates drag at all times.

**Fig 3.**
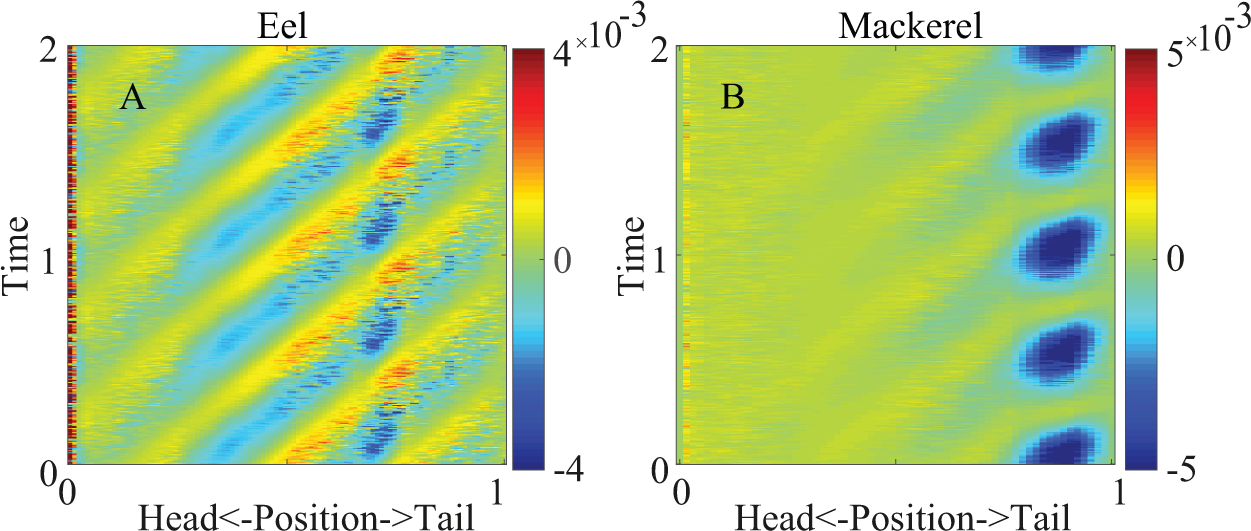
Spatiotemporal distribution of the fore-aft force (*F_x_*) on the eel (A) and the mackerel (B) for two periods. Negative values indicate thrust, as the swimming direction is in the -*x* direction.

**Fig 4.**
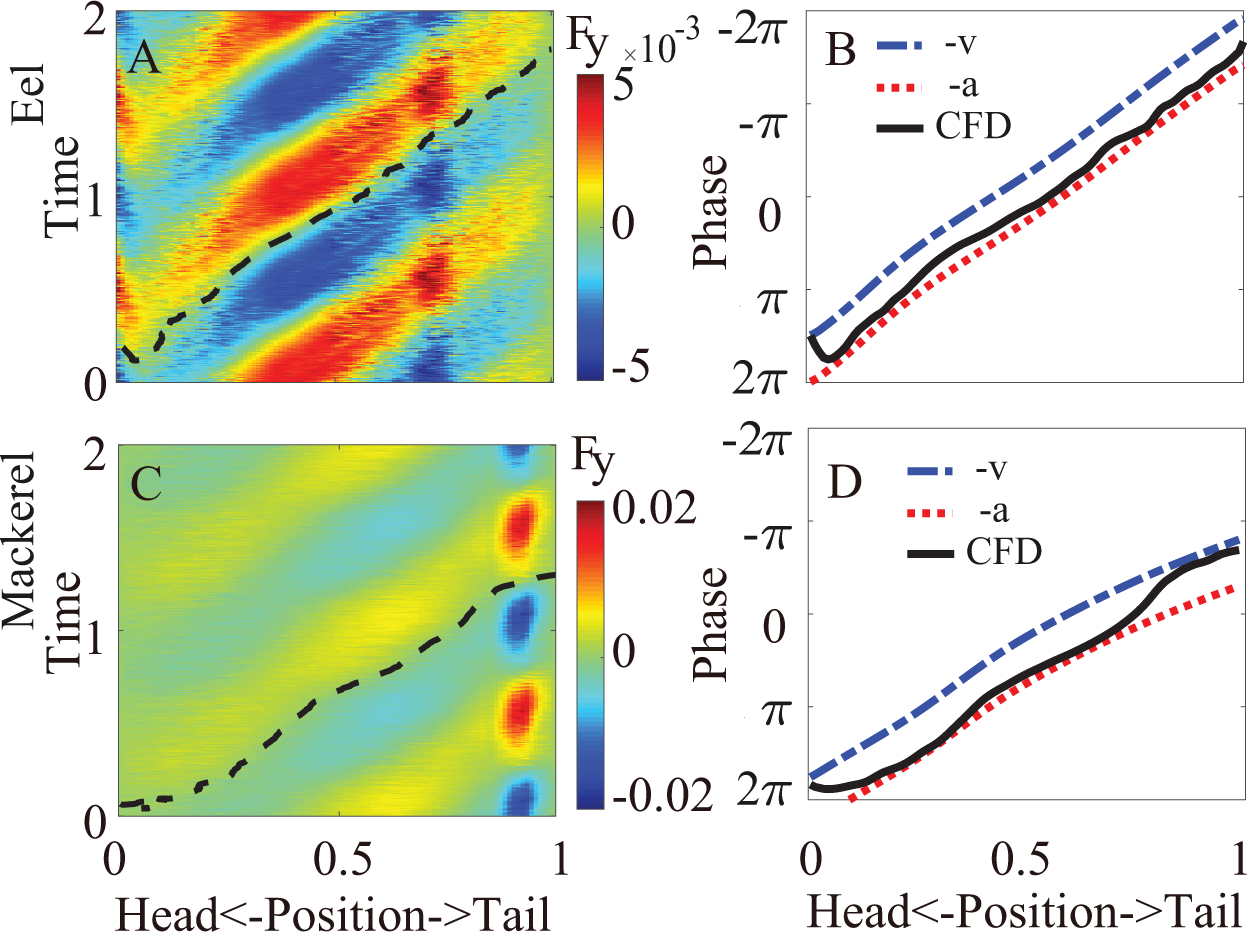
Lateral force (*F_y_*). Left column: Spatiotemporal distribution of the lateral force on the body for two periods from the simulation. The dashed black line indicates a zero-crossing (phase) of the force. Right column: Comparison of the phase of the lateral force along the body from CFD (solid black line, the same as the dashed lines in A and C) with the phase of the negation of the velocity and the phase of the negation of the acceleration. A 2*π* term is added or subtracted to ensure continuity.

Because the phase of the force, especially the lateral force, is essential in determining the phase of the torque [18], we compare the phases of the lateral forces from the simulation with those of the velocity and the acceleration of the segments. Roughly speaking, if the lateral force from the fluid on a segment is in phase with the negation of the segment velocity, it is a resistive-like force, and if the lateral force is in phase with the negation of the acceleration of the segment, it is a reactive-like force. We find that the phase of the observed lateral force on the body is closer to the phase of the negation of the acceleration except near the snout tips and the tail for the mackerel. In these regions, the phase of the lateral force is close to the negation of the velocity. Roughly speaking, the forces on the fish are close to the predicted forces from elongated body theory, but there are discrepancies when the shape changes are rapid. Detailed discussions of the hydrodynamics underlying the force pattern are beyond the scope of this paper.

### Torque

The torque required to overcome the hydrodynamic forces and body inertia in both species exhibits a traveling wave pattern moving posteriorly with a higher speed than the curvature wave (Fig 5). For the eel, the average speed of the torque wave (*v_T_*) is 1.43 in the nondimensionalized unit (body length/period), 2.4 times that of the curvature wave (*v_κ_* = 0.59). The traveling wave speed of the torque is even higher in the mackerel (*v_T_* = 2.23), exhibiting a nearly standing wave pattern. The torque wave speeds qualitatively match the observation that the EMG speed is much higher in carangiform swimmers (Fig 1). The maximal value of the torque appears at approximately the middle of the body of the eel and slightly posterior to the middle point for the mackerel.

**Fig 5.**
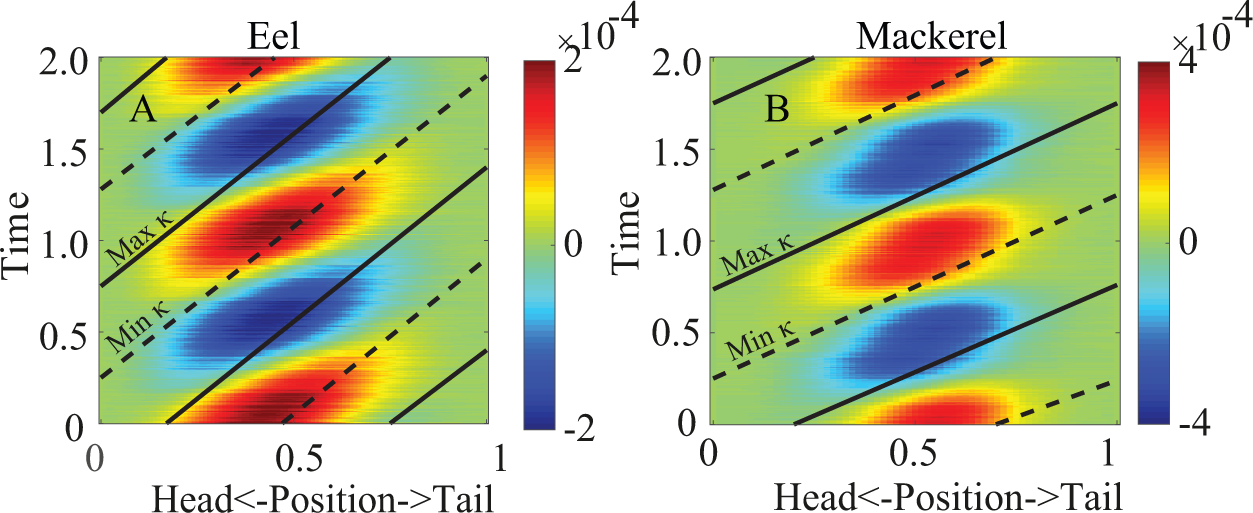
Spatiotemporal distribution of the torque on the body in two periods for the eel (A) and mackerel (B). The solid and dashed lines indicate the maximum and minimum curvatures, respectively. The same information is illustrated by S3 Video and S4 Video.

### Power

As shown in Fig 6, the power from the torque is mostly positive, indicating the energy output from the muscle, but negative values are observed on the posterior parts of both fish. For the eel, the power is nearly all negative for *s >* 0.6, similar to the case with a floppy body in a previous 2D study [12], while for the mackerel, the negative power is intermittent on the posterior part. The work over a cycle calculated by simply integrating the power is the minimal work needed, since the dissipation due to the internal resistance is not included; this method implies that the negative power transferred to the body is fully stored and recovered. The peak of this work per cycle is at the anterior part (≈0.4) for the eel and at a more posterior position for the mackerel (≈0.65), slightly posterior to the peak magnitude of the torque. We find that the work over a cycle is significantly negative on the posterior half of the eel body and slightly negative near the tail of the mackerel. If we assume that no energy-storing and transmitting elements exist, the work done by the muscles is the integration of only the positive power. We denote this quantity by *W* ^+^. The differences between the two kinds of work per cycle are the greatest for the posterior part of the eel, indicating that power is lost if no spatial energy transfer is performed inside the eel body. The distribution of power transferred to the fluid from the body is relatively uniform on the eel but concentrated on the tail of the mackerel (cyan dashed lines in Fig 6B&D).

**Fig 6.**
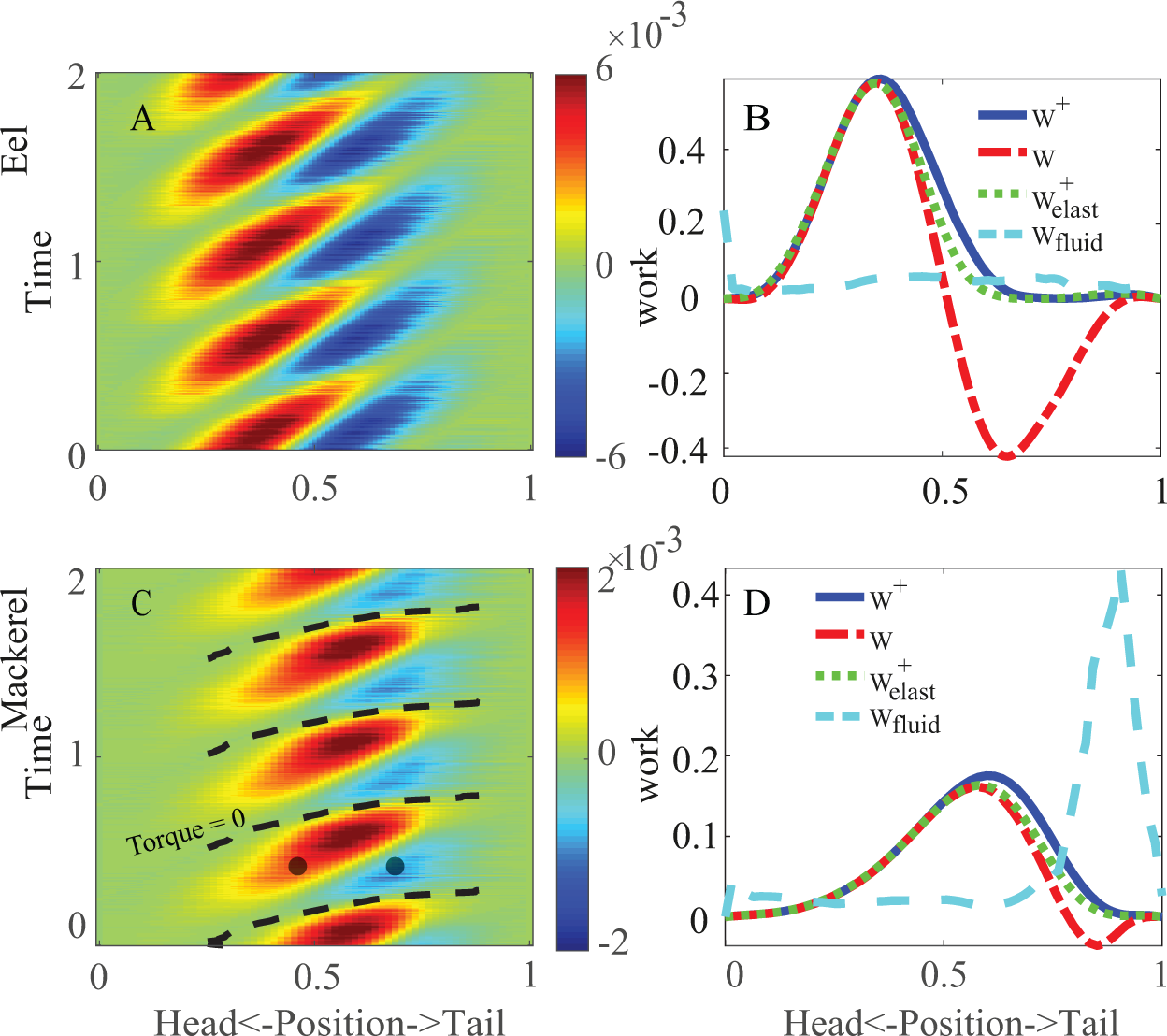
Internal power (*P_T_* (*s, t*)) distribution for the eel (A) and the mackerel (C) and the work done over a cycle by muscles along the body of the eel (B) and the mackerel (D). The dashed line in (C) indicates the zero-crossing of the torque in the mackerel (Fig 5B). The two black dots in (C) indicate an example time instant when two points on the body have the opposite sign of power but the same sign of torque. The solid blue lines in (B & D) represent the work by integrating only the positive values in (A & C), and the dashed-dotted red lines represent the work (*W*) by integrating both positive and negative values. The dotted green lines represent the positive work *W* ^+^ when body elasticity is considered (Fig 8B & D). The cyan dashed lines represent the work done to the fluid by the integration of *P_F_* (*s, t*).

The mean total power *P*_tot_ averaged over a cycle is 2.0 × 10^*−*4^ (in nondimensionalized units) for the eel and 2.5 × 10^*−*4^ for the mackerel. If only the positive power is used, the power becomes 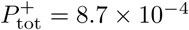 and 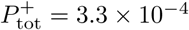, for the eel and the mackerel, respectively. The significant differences between *P*_tot_ and 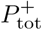 indicate the great potential to improve energetic efficiency by the spatiotemporal transfer of energy.

### Understanding the torque and power patterns

The torque pattern can be understood by applying the results obtained in a previous study [18]: The torque pattern in undulatory locomotion is determined mainly by the wavelength and phase of the lateral force relative to the lateral movement. The torque wave of the eel has a relatively low wave speed relative to that in the case of the mackerel due to the short wavelength of the undulation. Since the phase of the force for the eel is overall close to the phase of the reactive force, the internal torque and power patterns are also similar to the patterns associated with pure reactive forces (Fig 7, left column). For the mackerel, the long wavelength of the curvature wave and the concentrated force on the tail result in nearly synchronized torques on the body. Because the force from the tail to the fluid is nearly in phase with the velocity, the rate of change of curvature (**) and the torque are also nearly in phase. As such, the torque and power patterns are similar to the patterns associated with pure resistive forces (Fig 7, right column), and the internal power is nearly all positive.

**Fig 7.**
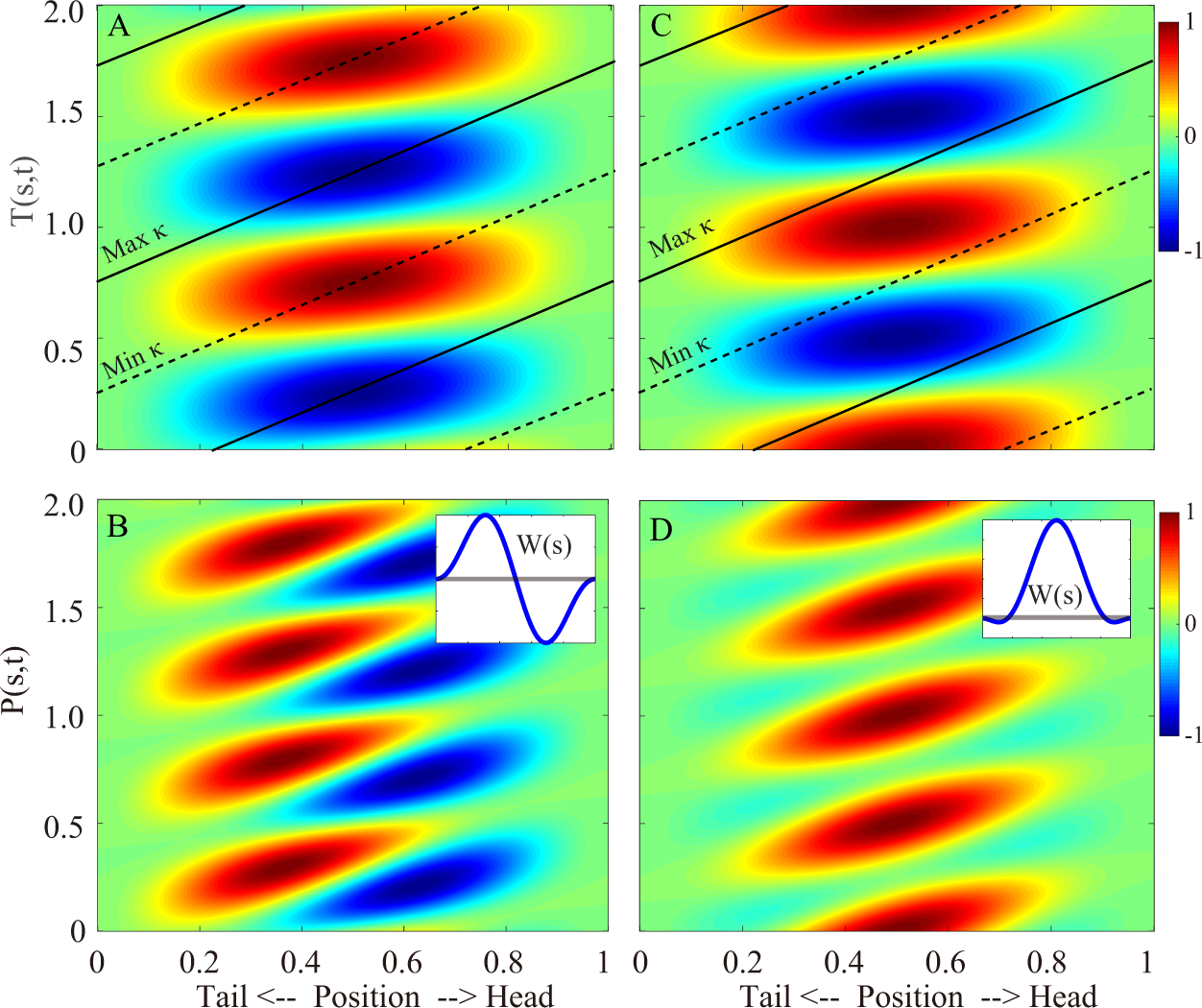
Spatiotemporal distributions of the torque (*T*) and power (*P*) along the body when pure resistive forces (left column) or pure reactive forces (right column) are considered. All values are normalized to the respective maximum values in each subfigure. The solid and dashed lines indicate the maximum and minimum curvatures, respectively. The insets are the distribution of the work 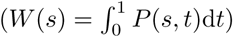 done by the internal torque. The gray lines in the insets indicate 0 to guide the eye. In the calculation, the body is uniform, the undulation amplitude is uniform and infinitesimal, and the wavelength is the same as the body length. See S2 Appendix for details of the derivation.

### Body viscoelasticity explains the wave speed variation of EMG

Previous bending tests and experiences in the handling of fish indicate that the torque from the viscoelasticity of an eel body is significant but smaller than the torque generated by muscles [19]. For carangiform swimmers, since no muscles exist behind the peduncle region and the curvature is comparable (albeit greater) with the rest of the body, the torque from elasticity must be significant at least in the tail region. However, an accurate *in vivo* measurement of the body viscoelasticity distribution is not available. Therefore, here we discuss the trend of the influences of the viscosity and elasticity individually when the elasticity or viscosity is small relative to the torque from hydrodynamics and the body inertia (Fig 8).

**Fig 8.**
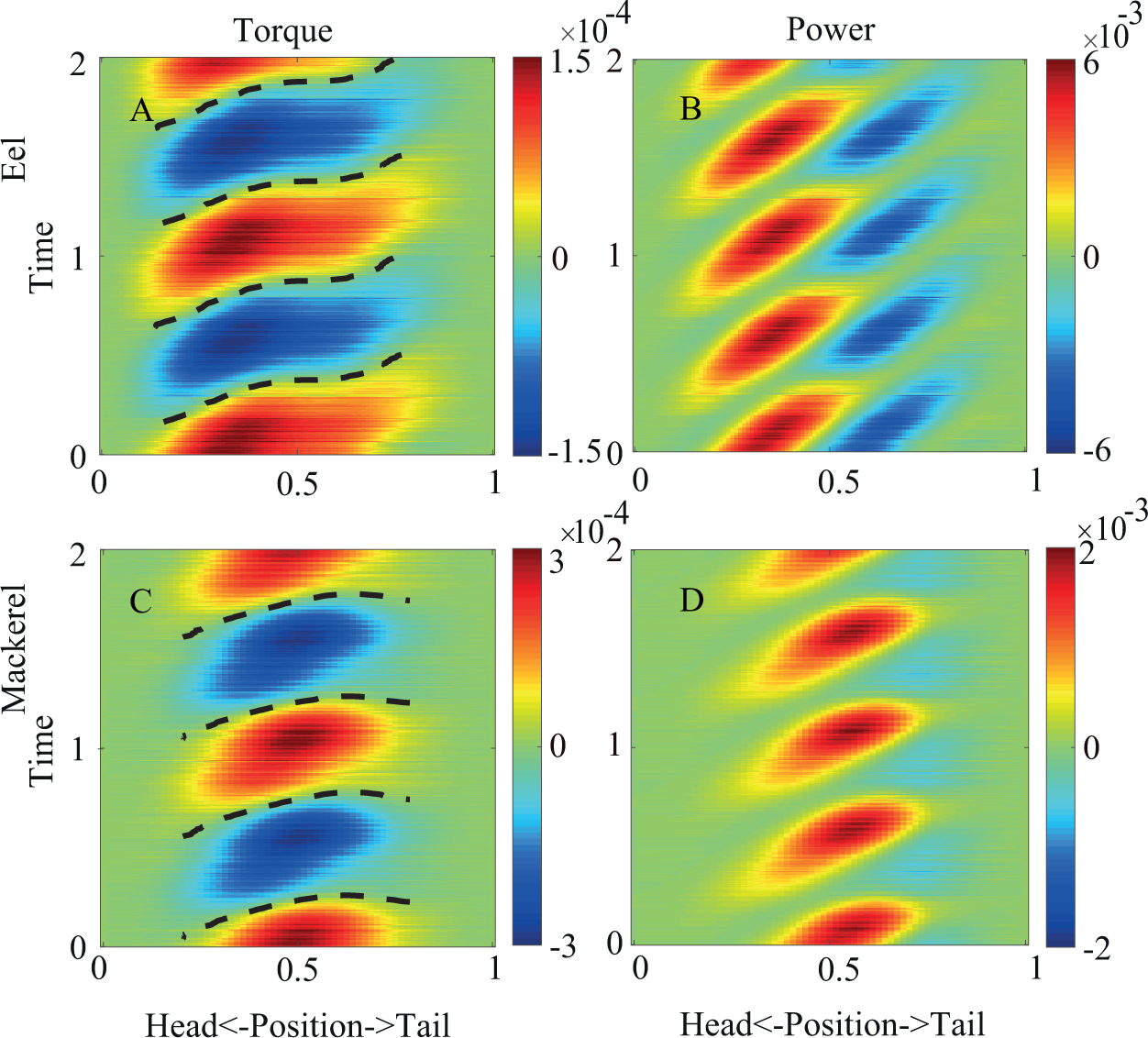
Torque (left column) and power (right column) distributions when the elasticity of the body is considered for the eel (top row) and the mackerel (bottom row). The dashed lines indicate zero-crossings of the torque.

**Fig 9.**
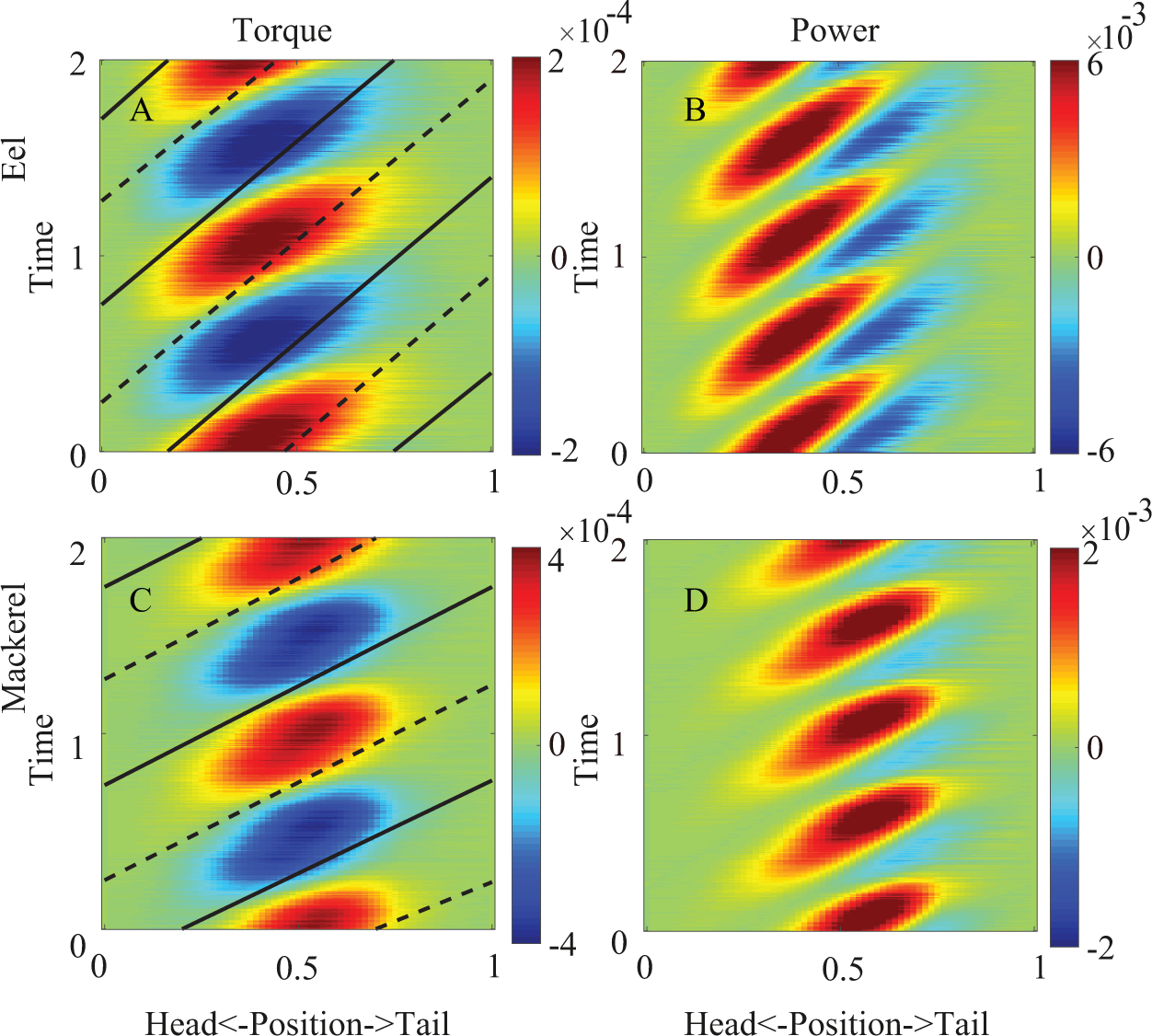
Torque (left column) and power (right column) distributions when the viscosity of the body is considered for the eel (top row) and the mackerel (bottom row). The solid and dashed lines indicate the maximum and minimum curvatures, respectively.

We assume that the magnitude of the torque from the body elasticity or viscosity is 20% of the torque at individual positions along the body, namely *T_e_*=0.2〈*T*〉*κ*(*s,t*)*/〈κ〉*or 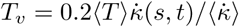, where “〈〉” means standard deviation over time. As shown in Fig 8, we find that the effect of elasticity on the torque is different along the body, separated by a position (*s* ≈0.5 for the mackerel and *s* ≈ 0.2 for the eel) where *T* and ** are in phase and the power is all positive. Anterior to that point, the torque magnitudes increase, and the torque wave speeds decrease; posterior to that point, the torque magnitude decreases, and the speed of the torque wave increases. For the mackerel, the torque wave can even reverse when the phase shift effect of the elasticity is strong. The reversal of the wave resembles the reversal of the wave of the offset of the EMG observed for carangiform swimmers (Fig 1). As a result of the changes in the torque, the area of the negative power region in the posterior part of the body decreases, and *W* ^+^ decreases (Fig 6D). This observation is consistent with the findings of previous studies that suitable elasticity can save and restore energy to improve efficiency (e.g., [20]). For the eel, the effect of the speed increase ends near *s* = 0.7 when the maximal curvature coincides with the minimal torque without elasticity. Therefore, the energy storing and releasing for the eel is in the middle part of the body (Fig 6B). Since the body viscoelasticity of the eel is weak and this effect is subtle, changes in the wave speed of the middle part of the torque wave or EMG are not obvious.

Since the body viscosity requires a torque in phase with the time derivative of the curvature, for both the eel and mackerel the resulted torques become more aligned with the time derivative of the curvature and hence have wave speeds closer to the speed of curvature (*v_T_ /v_κ_* = 2.2 for the eel and *v_T_ /v_κ_* = 1.8 for the mackerel). Consequently, the negative power regions are reduced since the viscosity of the body always dissipates energy.

### Tendon connection explains the duration decrease of EMG

While local elasticity can transfer energy temporally, the spatial transmission of energy can only be achieved by other structures. In animals, coupled joint articulation by tendons over two or more joints is common and is an effective structure to save and transfer energy [21]. For carangiform swimmers, long tendons exist that span over many vertebra [22]. We hypothesize that these long tendons are used to transfer energy from the posterior part to the middle part of the body when the negative power appears on the posterior part (Fig 6). This hypothesis can explain the observed decrease in the muscle activation duration among the carangiform swimmers, including some detailed features: the increase in the duration of the negative power from the middle of the body towards the tail matches the decrease in the EMG duration. The start of the positive power is aligned with the sign change of ** (the lines in Fig 5B), resulting in a low speed that is the same as that of the curvature wave. The end of the positive power is aligned with the sign change in the torque (the dashed lines in Fig 6B), resulting in a high speed that is the same as that of the torque wave. Such differences in wave speed qualitatively match the speed differences of the onset and offset of the EMG. Note that this hypothesis does not contradict the common view that force and energy are transmitted to the tail to interact with the fluid. Actually, the torque is still required when the power is negative on the posterior region and can be provided by the muscle in a more anterior position connected by the tendon. This hypothesis is also consistent with the observation that the EMG duration is nearly half of the undulation period on the whole body of anguilliform swimmers, which do not possess long tendons [22].

### Error associated with *Re*

The low swimming speeds we observed (compared with those of real animals) are likely due to the low *Re* used in our simulations. However, we argue that the results are qualitatively representative for real adult fish. First, a meta-analysis of previously reported fish swimming data indicates that the transition from the viscous regime to the turbulent regime occurs at a *Re* of several thousand [23]. Second, even the eel model in our study shows an inertia-dominated mode of swimming. Since the drag coefficient decreases with increasing *Re* in general, the speed of the simulated swimmer is expected to increase with increasing *Re*, and the contribution of the resistive force is expected to decrease for a real adult eel.

## Conclusion

Using 3D numerical models, we provide so far the most accurate prediction of the torque and power required for hydrodynamic forces during the undulatory swimming of fish. By considering the torque and power transfer by tendons and the body viscoelasticity, we for the first time give explanations for some long-standing questions in muscle activation patterns. Our study offers an integrative view of the function of the muscles as part of the mechanical system, highlights the differences in the internal driving of two primary swimming modes, and provides insights on the energy transfer and saving mechanisms of fish swimming. The numerical models developed and the mechanisms revealed in this study may guide the design of efficient bio-inspired robots, especially soft robots with distributed driving systems and elastic bodies [24, 25].

## Supporting information

**S1 Appendix. Description of the method to satisfy momentum conservations and obtain kinematics in the body frames.**

**S2 Appendix. Derivation of the torque and power from pure resistive forces and reactive forces.**

**S1 Fig Mesh used in the simulation.** (A) Eel. (B) Mackerel.

**S2 Fig The kinematics of the swimmers in their own body frames (free movement in the vacuum)** (A) Eel. (B) Mackerel.

**S1 Video. The force distribution along the body of the eel.** The black line represents the midline of the fish and the red arrows represent the hydrodynamic forces. The head is on the left and the tail is on the right.

**S2 Video. The force distribution along the body of the mackerel.** The black line represents the midline of the fish and the red arrows represent the hydrodynamic forces. The head is on the left and the tail is on the right.

**S3 Video. The torque distribution along the body of the eel.** The magnitude of the torque in the z direction is represented by the color.

**S4 Video. The torque distribution along the body of the mackerel.** The magnitude of the torque in the z direction is represented by the color.

## Acknowledgments

This work was supported by the National Natural Science Foundation of China grant No. 11672029 and NSAF-NSFC grant No. U1530401 (both to T.Y.M., B.W.J., J.L.S. and Y.D.). We thank Prof. Fotis Sotiropoulos and Prof. Iman Borazjani for sharing the shape data of the fish.

